# ELR is a true pattern recognition receptor that associates with elicitins from diverse *Phytophthora* species

**DOI:** 10.1101/2020.09.21.305813

**Authors:** Emmanouil Domazakis, Doret Wouters, Jan Lochman, Richard G. F. Visser, Matthieu H. A. J. Joosten, Vivianne G. A. A. Vleeshouwers

## Abstract

The first layer of plant immunity against pathogens is mediated by cell surface pattern recognition receptors (PRRs) that recognize pathogen molecules in the apoplast. Several pairs of PRRs and their matching extracellular ligands have been described but, in many cases, actual evidence for ligand binding by the PRR is lacking. The receptor-like protein ELR from *Solanum microdontum*, which triggers cell death upon co-expression with elicitins of various *Phytophthora* species and enhances resistance to late blight caused by *Phytophthora infestans*, was previously identified as the elicitin receptor by forward genetic screenings employing the INF1 elicitin of *P. infestans*. In this study, we investigated whether ELR associates with INF1 and other elicitins that are secreted by diverse *Phytophthora* spp. We performed *in planta* and *in vitro* co-immunoprecipitation of ELR with several affinity-tagged elicitins, as well as *in planta* transient co-expression assays. We found that ELR physically interacts with the class I elicitins INF1 and ParA1, from *P. infestans* and *Phytophthora parasitica*, respectively, which is in line with their ability to cause cell death when co-expressed with ELR in potato. Together, we demonstrate that ELR is a genuine PRR that binds elicitins of *Phytophthora* species.

## Introduction

A major part of a plant’s defense response against microbes depends on the effective detection of immunogenic patterns, such as microbe-associated molecular patterns (MAMPs) (Couto and Zipfel, 2016; Gust and Felix, 2014; Jones and Dangl, 2006; Ranf, 2017; van der Burgh and Joosten, 2019). MAMPs are recognized in the plant apoplast, typically by membrane-localized pattern recognition receptors (PRRs). All plant PRRs that have been identified so far contain a variable number of leucine-rich repeats (LRR), lysine motifs or lectin domains in their ecto-domain, a transmembrane domain and may contain a kinase domain. LRR-type receptor-like proteins (RLPs) and receptor-like kinases (RLKs) predominantly recognize proteinaceous MAMPs or effectors, of which the latter are compounds produced by pathogens with the aim to perturb plant defense responses. Despite the high amount of PRRs encoded by plant genomes, and the increasing number of PRRs discovered over the past 20 years, for most of them it has not been determined whether they are actually indeed true receptors of their matching ligands, or just involved in the complex that mediates resistance to pathogens (Boutrot and Zipfel, 2017; Dodds and Rathjen, 2010; Tang et al., 2017; Zipfel, 2014). RLPs are relatively understudied when compared to RLKs, and providing evidence for binding of MAMPs or effectors has been challenging and is rather limited. Successful examples of RLPs for which ligand binding has been shown are the *Arabidopsis* RLP42/RBPG1 and RLP23, which associate with a fungal endo-polygalacturonase and the widely conserved MAMP nlp20, respectively (Albert et al., 2015; Zhang et al., 2014). From Solanaceous plants, LeEIX2 from tomato was found to associate with an ethylene-induced xylanase (Ron and Avni, 2004) and the LRR-RLP RXEG1 from *Nicotiana benthamiana* binds the glycoside hydrolase PsXEG1 from *Phytophthora sojae* and provides resistance to this oomycete pathogen (Wang et al., 2018).

Elicitins are a family of structurally conserved extracellular oomycete proteins, specific to the genera *Phytophthora* and *Pythium*. Elicitins have features of oomycete MAMPs, as they have been shown to induce localized cell death, salicylic acid-, jasmonic acid- and ethylene-dependent accumulation, as well as systemic resistance in several responding plant genotypes, across various families (Derevnina et al., 2016; Vleeshouwers et al., 2006). The Solanaceous plants *Nicotiana benthamiana, N. tabacum* and *S. microdontum* mount a specific cell death response upon treatment with the INF1 elicitin (Du et al., 2015; Huitema et al., 2005; Kamoun et al., 1998; Vleeshouwers et al., 2006). All elicitins share a highly conserved 98-amino-acid elicitin domain, which contains six cysteine residues at conserved positions for the formation of disulfide bridges (Derevnina et al., 2016; Jiang et al., 2006). Elicitins of the canonical class ELI-1 consist of a signal peptide for extracellular targeting, which is cleaved off upon secretion, and the elicitin domain only. Elicitins of all other classes possess an additional C-terminal extension, which is of variable length and is putatively involved in plasma membrane or cell wall anchoring.

ELR, an RLP from the wild potato species *Solanum microdontum*, mediates recognition of various elicitins of diverse *Phytophthora* species (Du et al., 2015). The recognition spectrum ranges from elicitins belonging to class ELI-1 (e.g. INF1, ParA1), ELI-2 (e.g. INF2A) and class ELI-4 (e.g. INF5, INF6) (Du et al., 2015; Jiang et al., 2006) (Figure S1). In contrast to bacterial flagellin, EF-Tu and NLPs (Bohm et al., 2014; Felix et al., 1999; Kunze et al., 2004), elicitins do not seem to share a conserved stretch of amino acids (Fig S1a), but rather the conserved overall structure of all elicitins is likely recognized by the extracellular LRRs of ELR (Derevnina et al., 2016). In line with this, the elicitins that are recognized by ELR share rather low amino acid similarities (Du et al., 2015) (Figure S1b).

ELR lacks a cytoplasmic kinase domain and is thus incapable of downstream signaling by itself. However, recently, it was shown that ELR physically associates with the RLK SUPPRESSOR OF BIR1-1 (SOBIR1) in a ligand-independent manner, and associates with the RLK SOMATIC EMBRYOGENESIS RECEPTOR KINASE 3 (SERK3), also referred to BRI1-ASSOCIATED KINASE 1 (BAK1), upon ELR activation by elicitins (Domazakis et al., 2018; Du et al., 2015). Both SOBIR1 and SERK3 are required for ELR function and for triggering resistance to pathogens, including *Phytophthora infestans* (Chaparro-Garcia et al., 2011; Domazakis et al., 2018; Liebrand et al., 2014; Peng et al., 2015).

In this study, we provide evidence that ELR physically associates with INF1 elicitin, using *in planta* and *in vitro* approaches. Furthermore, we tested various additional elicitins of different *Phytophthora* species and found a correlation between physical interaction of the elicitin with ELR and the occurrence of an ELR-mediated cell death response. Our data render ELR a genuine PRR, which associates with various members of the elicitin family of *Phytophthora* species.

## Materials and methods

### Plant materials

*Nicotiana benthamiana* plants were grown from seeds and *Solanum hjertingii* genotype 349-3 (HJT349-3) plants were clonally propagated *in vitro* on Murashige and Skoog (MS) medium, supplemented with 20% w/v sucrose as described before (Du et al., 2015; Murashige and Skoog, 1962). Seedlings of *N. benthamiana*, as well as well-rooted HJT349-3 plantlets, were transferred to pots with disinfected soil in a climate-regulated greenhouse compartment at Unifarm, Wageningen University & Research (22/18°C and 16/8 h light day/night regime at 70% relative humidity) where they were grown for 3 weeks prior to agroinfiltration experiments. The transient protein expression and cell death assays were performed under the same conditions.

### Binary vectors for *Agrobacterium tumefaciens*-mediated transient transformation

Construction of pBin-KS-*p35s:ELR-eGFP*, pBin-KS-*p35s:Cf-4*-*eGFP*, pBin-KS-*p35s:eGFP* and pCB302-3-*p35s:INF1* has been described earlier (Du et al., 2015; Liebrand et al., 2013). Full length elicitin constructs of INF1, INF1^C23S^, ParA1, CRY2, INF2A, INF2B, INL1 and PYU1were synthesized, with their native signal peptide (Sp) replaced with the pathogenesis-related protein (PR1) Sp from tobacco as well as two N-terminal HA-tags, separated by spacers (Figs. S1a, S2a), as previously described (Domazakis et al., 2017). The PR1Sp2HA elicitin sequences were flanked by AttL1 and AttL2 recombination sites to facilitate direct Gateway recombination cloning (Invitrogen). The constructs were synthesized in pUC57 vector (GenScript). From those pUC57 constructs, the elicitin domains of INF2A (amino acid (aa) 21-118), INF2B (aa 21-118), CRY2 (aa 21-118) and INL1 (aa 19-104) (elicitin domains were defined according to (Jiang et al., 2006)) were PCR-amplified using Phusion DNA polymerase (Thermo Fisher Scientific) and primers listed in Table S1, and were cloned into pENTR/D-TOPO (Invitrogen) (Figs. S1a, S2a). Then, together with INF1, INF1^C23S^, ParA1 and PYU1, they were recombined into the pK7WG2 binary vector by a Gateway LR reaction (Invitrogen). Final constructs were transformed into *A. tumefaciens* stain AGL1.

### *Agrobacterium* transient cell death assays

For elicitin cell death induction assays, young, fully expanded leaves of 3 week-old *N. benthamiana* or *N. tabacum* SR1 plants, were agroinfiltrated with pK7WG2-empty vector (EV), pCB132:*p35s-INF1* and pK7WG2:*p35s*-*PR1Sp2HA-elicitins* constructs, at an OD_600_ of 0.2. Similarly, *Solanum hjertingii* 349-3 plants were co-infiltrated with 1:1 mixtures of *A. tumefaciens* carrying pK7WG2:*p35s-ELR* (OD_600_=0.15), in combination with pK7WG2-EV, pCB132:*p35s-INF1* or pK7WG2:*p35s*-*PR1Sp2HA-elicitins* constructs (OD_600_=0.1). Leaves were visually examined for cell-death at 3-5 dpi. Cell death quantifications were performed as described previously (Du and Vleeshouwers, 2014).

### Co-immunoprecipitations and immunoblotting

*In planta* protein-protein interaction studies were performed by transient expression of both proteins of interest in young fully expanded leaves of 3 week-old *N. benthamiana* plants using *Agrobacterium*-mediated transformation, followed by co-immunoprecipitation. At 2 days post infiltration, leaves were collected, snap-frozen in liquid nitrogen and ground to a fine powder. Protein extraction, co-immunoprecipitation and western blot analysis with αGFP-HRP and αHA-HRP was performed as described (Domazakis et al., 2018). Immunoprecipitations were performed using GFP-trap_MA (50% slurry) (Chromotek) and Pierce Anti-HA magnetic beads (Thermo Fisher Scientific).

### Production and purification of recombinant proteins

Recombinant His-HA-INF1 and the unrelated His-HA-SCR74, a 74 amino acid cysteine-rich protein from *P. infestans* with similarity to the PcF protein from *Phytophthora cactorum fragaria* (variant B3b, NCBI accession number AY723717.1), were synthesized in a codon-optimized version for expression in *Pichia pastoris* (GenScript). Codon-optimized coding regions were designed to carry N-terminal His-HA tags, as described (Fig. S2b) (Domazakis et al., 2017). The coding regions were synthesized with flanking *Stu*I and *Kpn*I restriction sites to facilitate cloning and were obtained in the pUC57 vector. Coding regions were cloned in frame with the α-mating factor in the pPinkα-HC vector (Invitrogen), enabling protein secretion. The *P. pastoris* PichiaPink™ strain 1 (Invitrogen), was transformed by electroporation and high protein producing clones were selected using a high throughput screening method (Domazakis et al., 2017). For protein production, yeast cultures were first grown in 1 L of buffered glycerol complex medium (BMGY), at 28°C, and shaking at 250 rpm, till an OD_600_ of 2-6 in a 3 L conical flask. Cells were then pelleted by centrifugation and were resuspended in 200 mL of buffered methanol complex medium (BMMY), containing 1% v/v MetOH at 28°C, and shaking at 300 rpm, for induction of protein production in a 2 L conical flask. Induction conditions were maintained for 48 hours with extra MetOH added to maintain 1% v/v, every 24 hrs. After 48 hours, the cultures were harvested, and the cells were removed by centrifugation. Supernatants were filter-sterilized through a 0.22 μm syringe filter. A pressurized Amicon stirred cell concentrator (Millipore), combined with a 3 kDa exclusion filter, was used to concentrate the supernatants to a volume of ∼10 mL. Purification of the expressed protein was performed using cation exchange. For this, a column was assembled containing SP Sepharose, for which the beads were equilibrated with 10 volumes of acetic acid buffer (50 mM acetic acid, pH 4.5). Concentrated yeast supernatants containing elicitins were diluted five times in the acetic acid buffer and were slowly loaded onto the column. The column was then washed with five volumes of the acetic acid buffer, and proteins were subsequently eluted by applying five volumes of elution buffer (10 mM Tris, 500 mM NaCl, pH 8) and collecting the eluate in several fractions. The protein-containing fractions were obtained by measuring A280 absorbance, SDS-PAGE and western blotting. Protein concentrations were determined using a BCA assay (Thermo Fisher Scientific).

### *In vitro* binding assay

*A. tumefaciens* strains carrying ELR-eGFP or Cf-4-eGFP were infiltrated into leaflets of *N. benthamiana* plants at an OD_600_ of 1 and 0.8, respectively. Two days after agroinfiltration, leaves were collected and snap-frozen in liquid nitrogen. Leaves were ground to a fine powder and RIPA extraction buffer was added to an amount of 2 mL/g of leaf material (Liebrand et al., 2012; Liebrand et al., 2013). Purified recombinant HisHA-INF1 or HisHA-Scr74 were incubated with 20 μl Pierce Anti-HA magnetic beads (Thermo Fisher Scientific) at an amount of 500 pmol each for 30 min. Subsequently, beads were washed four times with binding buffer (100 mM MES, 10 mM NaCl, 3 mM MgCl_2_, pH=6 (Saur et al., 2016)). Then the extract containing ELR-eGFP or Cf4-eGFP was diluted at a ratio of 1:3 in the same binding buffer and 2 mL of that extract was applied to His-HA-INF1- or His-HA-SCR74-bound beads. The mixture was incubated for 1 hour at 4°C, after which co-immunoprecipitation was performed as described (Domazakis et al., 2017; Domazakis et al., 2018).

## Results and discussion

### ELR associates with INF1 *in planta*

To study whether ELR associates with INF1 in co-immunoprecipitation experiments, we generated an N-terminally tagged version of INF1. INF1 was designed with the tobacco pathogenesis-related protein 1 (PR1) signal peptide (Sp) for secretion, and a 2×HA tag (PR1Sp2HA-INF1) (Fig. S2a). To test whether ELR associates with INF1 *in planta*, we co-expressed PR1Sp2HA-INF1 with ELR-eGFP (Du et al., 2015), and used Cf-4-eGFP or eGFP only, as negative controls for ELR-eGFP. In addition, as another negative control for INF1, we included a cysteine mutant of INF1 (PR1Sp2HA-INF1^C23S^), lacking in its overall structure one of the three disulfide bridges and a suppressed cell death response (Kamoun et al., 1997). After performing a co-immunoprecipitation experiment, we found that ELR specifically immunopurifies with INF1 (Fig. 1a). As expected, Cf-4 and eGFP did not associate with INF1, whereas the INF1^C23S^ mutant did show a severely reduced interaction with ELR, which is in line with the reduced cell death phenotype of this INF1 mutant (Kamoun et al., 1997). Interestingly, we noticed that INF1 migrates as a double band in the total protein extract, while after immunoprecipitation of ELR, only a single band is found (Fig. 1a, inputs). We propose that the upper band of INF1 corresponds to the unprocessed version of the protein, while the lower band, representing INF1 that associates with ELR, corresponds to the mature INF1 protein from which the Sp has been cleaved off. Alternatively, INF1 could be modified when expressed *in planta*, as has been suggested for *P. cryptogea* β-CRY (Uhlikova et al., 2016). In conclusion, these data show that ELR specifically associates with INF1 *in planta* and that this interaction occurs with the mature form of the elicitin, likely in the plant apoplast.

**Fig. 1.**
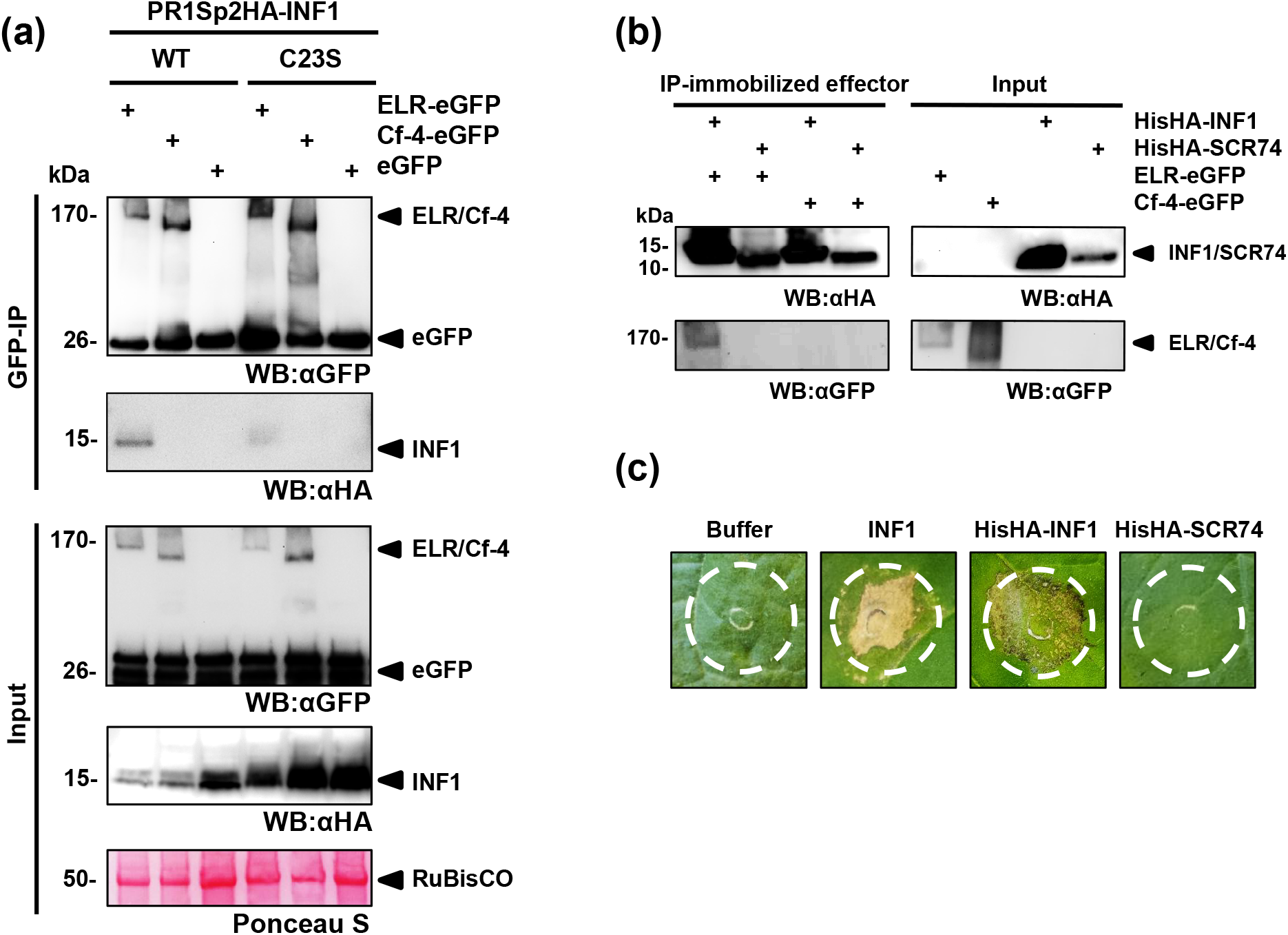
ELR associates with INF1 elicitin *in planta* and under *in vitro* conditions. (a) N-terminally HA-tagged INF1 wild type (PR1Sp2HA-INF1^WT^; WT) and the INF1^C23S^ cysteine mutant (PR1Sp2HA-INF1^C23S^; C23S) were co-expressed with ELR-eGFP, Cf-4-eGFP or eGFP only, in leaves of *Nicotiana benthamiana*. Total protein was extracted at 2 days post agro-infiltration and was subjected to immunoprecipitation using anti-GFP beads to capture ELR-eGFP, Cf-4-eGFP and eGFP. The immunopurified proteins were detected with anti-GFP, while their interaction with INF1 was assessed with anti-HA antibodies. Ponceau S staining of RuBisCO indicates equal protein loading. The results shown here are representative from three biological repeats. (b) Purified recombinant His-HA-INF1 or His-HA-SCR74 protein (500 pmol each) was immuno-absorbed to anti-HA beads for 30 min. Subsequently, beads were washed with binding buffer and diluted extract from leaves of transiently transformed *N. benthamiana* expressing ELR-eGFP or Cf-4-eGFP, was added to the beads (panel: input). After a 1h incubation, proteins were separated on SDS-PAGE gels and western blotting was performed. The immunopurified effector proteins His-HA-INF1 and His-HA-SCR74 were detected with anti-HA antibodies, while the interaction with the eGFP-tagged receptors was assessed with anti-GFP antibodies (panel: IP-immobilized effector). Results that are shown are representative for two biological repeats. (c) Functionality test of a 1 μM concentration of recombinant His-HA-INF1 and His-HA-SCR74 proteins in *N. benthamiana*, as compared to INF1 produced by *P. infestans*, at a similar concentration (all proteins were diluted in Milli-Q water).

### ELR associates with INF1 *in vitro*

To further validate the association between ELR and INF1, we designed an *in vitro* binding assay in which purified INF1 protein was tested for interaction with ELR. For this, His-HA-tagged INF1, as well as a His-HA tagged version of the unrelated protein SCR74, were produced in *Pichia pastoris* (Domazakis et al., 2017; Liu et al., 2005) and purified using cation exchange chromatography. To test for association, ELR-eGFP and Cf-4-eGFP were transiently expressed in *N. benthamiana* as described above, and were tested for interaction with the effector proteins. First, the His-HA-INF1 and His-HA-SCR74 proteins were immuno-absorbed to anti-HA beads. *N. benthamiana* extract, containing either ELR-eGFP or Cf-4-eGFP, was applied to the beads after three times dilution in binding buffer. After incubation at room temperature for 1 hour, and thorough washing with binding buffer we found that ELR-eGFP is specifically being captured from the plant extract by INF1, but not by the unrelated protein SCR74 (Fig. 1b). In contrast, as expected, Cf-4-eGFP was not interacting with either of these effectors. We confirmed that the recombinant His-HA-INF1 is recognized in *N. benthamiana* similar to untagged INF1, while His-HA-SCR74 does not trigger an HR Fig. 1c). These data show that ELR specifically interacts with INF1, and this finding is in accordance with our *in planta* observations.

### ELR associates *in planta* with ELI-1 elicitins

In contrast to other PRRs that recognize small epitopes of their matching MAMP and often do not trigger cell death, ELR seems to recognize some kind of structural domain of elicitins, and does induce cell death in plants (Derevnina et al., 2016; Du et al., 2015). To determine whether the cell death-inducing activity of the various elicitins is associated with their ability to bind to ELR, we generated affinity-tagged versions of diverse elicitin domains from various *Phytophthora* and one *Pythium* species. These included the N-terminally -HA-tagged elicitin domains of INF1, INF2A, INF2B and INL1 from *P. infestans*, ParA1 from *P. parasitica*, CRY2 from *P. cryptogea* and PYU1 from *Pythium ultimum*. The tested elicitins belong to different classes, INF1, INF2A, INF2B, ParA1 (classes ELI-1 and ELI-2), INL1 (class ELL) and PYU1, representing an outgroup (Fig. S1c), and share 23-94% amino acid identity to INF1 (Fig. S1b). To test the cell death-inducing activities of the various elicitins, agro-infiltration was performed in combination with ELR, in *S. hjertingii*, a wild *Solanum* genotype that does not respond to elicitins (Fig. 2a, b). We found that the tagged elicitins INF1, ParA1, CRY2 (belonging to the ELI-1 family) and INF2A (belonging to the ELI-2 family), induced cell death when co-expressed with ELR, at 3 days post-infiltration, similar to what was reported before (Du et al., 2015) (Fig. 2a, b). The INF1^C23S^ mutant, INF2B, INL1 and PYU1 did not trigger cell death (Fig. 2a, b).

**Fig. 2.**
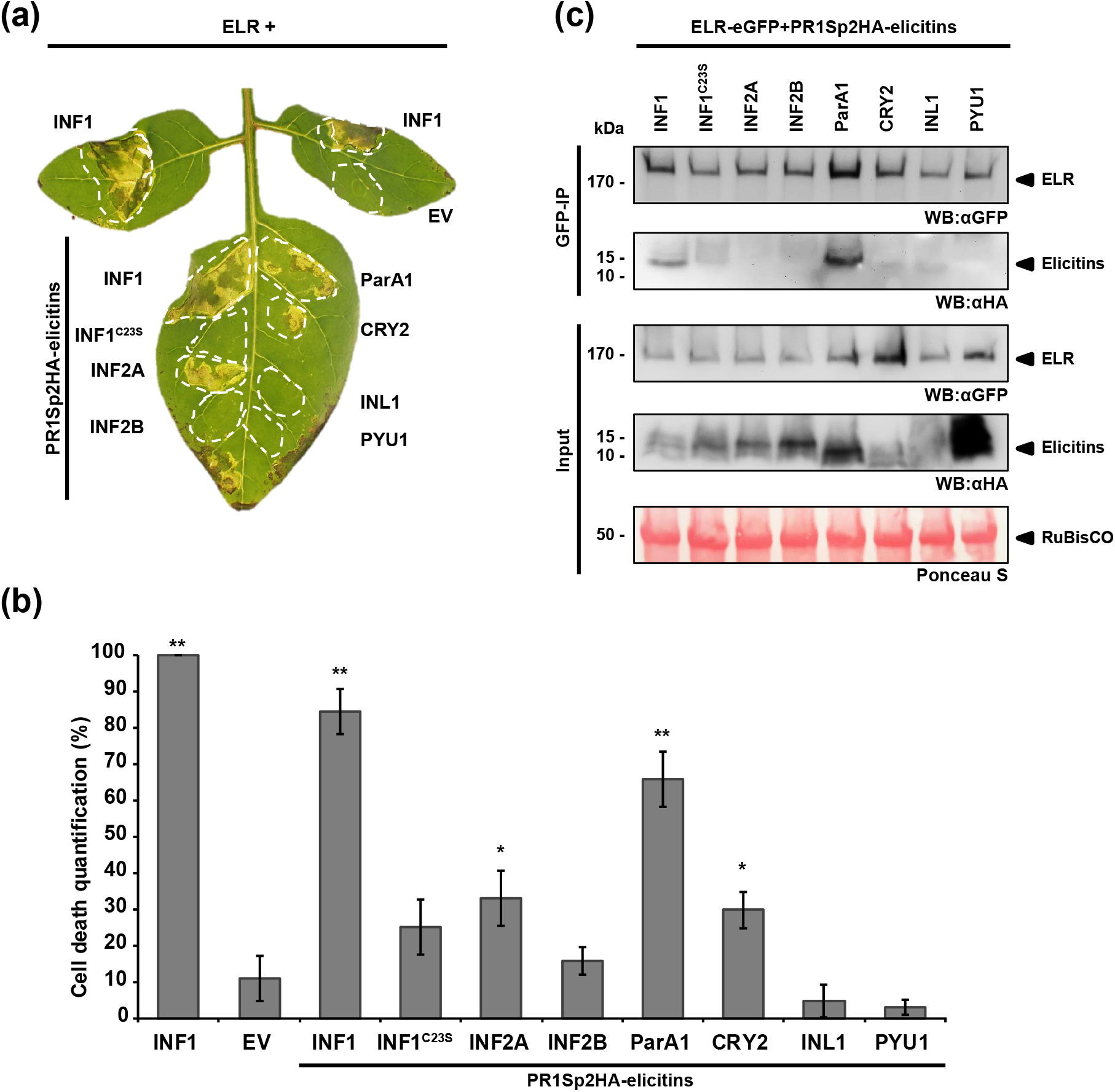
ELR associates with elicitins *in planta*. Diverse elicitins (ParA1, CRY2, INF2A, INF2B, and PYU1) from *P. parasitica, P. cryptogea, P. infestans* and *Pythium ultimum* as well as *P. infestans* elicitin-like INL1 were tested for recognition by ELR, together with INF1 and INF^C23S^ as a positive and a negative control, respectively. (a) N-terminally HA-tagged elicitin recognition in *Solanum hjertinjii* 349-5, upon their transient co-expression with ELR. Elicitin recognition by ELR results in cell death. (b) Quantification of the cell death response from the experiment shown in (a). (c) Co-immunoprecipitation of ELR with N-terminally HA-tagged elicitins. Total protein was extracted at 2 days post agro-infiltration of ELR in combination with the various elicitins, and was subjected to immunoprecipitation using anti-GFP beads, to capture ELR. Immunopurified ELR was detected with anti-GFP, while interaction with the elicitins was assessed with anti-HA antibodies. Ponceau S staining of RuBisCO indicates equal protein loading. Results that are shown are representative of three biological repeats.

To investigate whether the cell death-inducing activity of the various elicitins, correlates with their ability to associate with ELR, we performed an *in planta* co-immunoprecipitation (Co-IP) experiment of ELR-eGFP with the different affinity-tagged elicitins. N-terminally-HA-tagged INF1 and INF1^C23S^ were included as positive and negative controls, respectively (Fig. 2c). We found that, besides with INF1, ELR also interacts with ParA1, which belongs to the same ELI-1 class (Fig. S1c) (Fig. 2c).

For CRY2 and INF2A (Fig. S1a, b) that share lower amino acid identities to INF1 (74 % and 52%, respectively), we did not detect association with ELR (Fig. 2a, b). Since these elicitins do trigger cell death when co-expressed with ELR (Du et al., 2015), we anticipate that there should be interaction, however, we were unable to detect it in our Co-IP assay. Potentially this may be explained by reduced binding affinity of ELR to CRY2 and INF2A, in line with reduced cell death (Fig. 2b). INF2b, INL1 and PYU1 that do not cause cell death in agro-co-infiltrations with ELR, also do not show association with ELR. Overall, this study demonstrates that ELR is functioning as a genuine PRR and associates with elicitins beyond those produced by *P. infestans*, both *in vivo* and *in vitro*.

## Supplemental data

**Fig. S1. Characteristics of the elicitins that are subject of this study**. (a) The elicitin domains of seven elicitins from *Phytophthora* spp. and *Pythium ultimum* were aligned using COBALT, and BOXSHADE was used to visualize conservation. Amino acid residues identical in all sequences are highlighted in black, while those identical in >80% of the sequences are highlighted in grey. (b) Percentages of amino acid sequence identity of the elicitin domain between the different elicitins. (c) A neighbor-joining phylogenetic tree was created based on the elicitin domains of the seven elicitins using MEGA 5.1. Bootstrap support values (1,000 replicates) above 50% are given next to the branches.

**Fig. S2. Overview of affinity-tagged elicitins used in this study**. Effector constructs used for

(a) transient expression *in planta* using *A. tumefaciens* and for (b) recombinant protein expression in *Pichia pastoris*. Pathogenesis-related protein 1a (PR1a) or native INF1 signal peptides (Sp) for extracellular targeting were used, while elicitins were carrying hemagglutinin (HA) or histidine (His) tags, as indicated. The 98-amino acid elicitin domains were used for all tested elicitins.

**Table S1**. Primers used in this study.

## Acknowledgements

This work was supported by NWO-VIDI grant 12378 and COST Action SUSTAIN FA1208. We thank Sophien Kamoun for the always inspiring discussions, Prof. Dr. Evert Jacobsen for reviewing the manuscript, Isolde Bertram-Pereira for culturing *Solanum* plants, Henk Smid and Harm Wiegersma for help in the greenhouse.

