## Supplementary material for "ELR is a true pattern recognition receptor that associates with elicitins from diverse *Phytophthora* species": Supplemental data.pdf

**Supplemental data (Domazakis et al)**

**(a)**

```
INF1        1 ----TTCTTSQQTVAYVALVSILSDTSFNQCSTDSGYSMLTATSLPTTEQYKLMC
ParA1       1 ----TTCTTTQQTAAYVALVSILSDTSFNQCSTDSGYSMLTATSLPTTEQYKLMC
CRY2        1 ----TACTTTQQTAAYVALVSILSESFFSTCASDSGYSMLTATALPTTAQYELMC
INF2A       1 ----ETCSPTDQTTAYSTLASVLTLSSFQGCADDSGFSLLYSTALPDDDQYVKMC
INF2B       1 ----ETCSPTDQTTAYSTLASVLTLSSFQGCADDSGFSLLYSTALPDDAQYVKMC
INL1        1 ----ASCDVT-------SLQTLLTSSDTTTCATDSGYT-VTSLATPTDAEMDVMC
PYU1        1 YDEVTECPATE----FLKLAPLAANPNLSVCQDASGWQMLPPVGYPTDAQRALMC
consensus   1     ..*. . .. ... * .... .... *...**..... ...*..... .**

INF1       52 ASTACKTMINKIVSLNAPDCELTVPTSGLVLNVYSYANGFSSTCASL
ParA1      52 ASTACKTMINKIVTLNPPDCELTVPTSGLVLNVFTYANGFSSTCASL
CRY2       52 ASTACQEMIEEIIALNPPDCDLTVPTSGLVINVYEYANDFASTCASL
INF2A      52 ASDNCKSLIESVASLNPPNCDLTVPTSGLVLNVVDLTSGFSEKCSSS
INF2B      52 ASDNCKSLIESVAGLNPPNCDLTVPTSGLVLNVVDLTSGFSKKCSSS
INL1       44 TSTACQSVLSQLETLAPSECTL-----GTFALYADLITPLSNHCAGA
PYU1       52 VTPECFNLIDAIKALNVSDCVLVF--GDVKLNVKKLIEEFEPSCF--
consensus  56 ....*..... . .*....*.*........... ..  ... .*..
```

**(b)**

| Elicitin | INF1 | ParA1 | CRY2 | INF2A | INF2B | INL1 | PYU1 |
|---|---|---|---|---|---|---|---|
| **INF1** | ID | | | | | | |
| **ParA1** | 93.88 | ID | | | | | |
| **CRY2** | 74.49 | 76.53 | ID | | | | |
| **INF2A** | 52.04 | 53.06 | 52.04 | ID | | | |
| **INF2B** | 51.02 | 53.06 | 53.06 | 96.94 | ID | | |
| **INL1** | 27.06 | 29.41 | 32.94 | 30.59 | 31.76 | ID | |
| **PYU1** | 30.00 | 30.00 | 32.22 | 30.00 | 31.11 | 23.46 | ID |

**(c)**

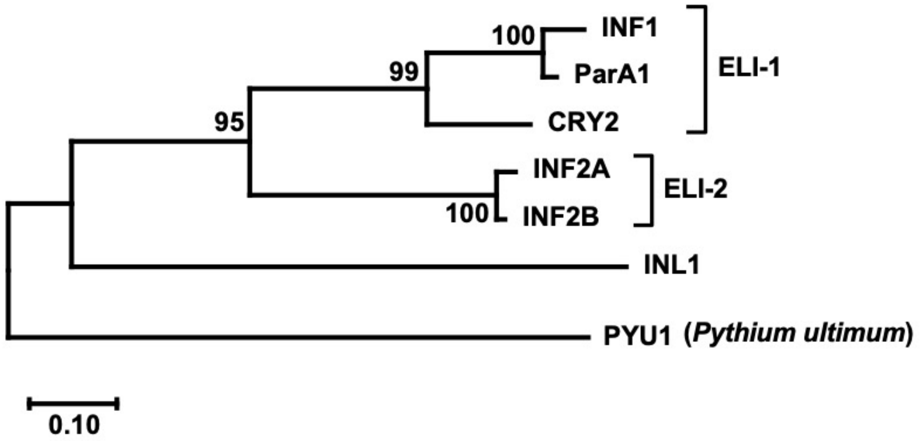

**Fig. S1. Characteristics of the elicitins that are subject of this study.** (a) The elicitin domains of seven elicitins from *Phytophthora* spp. and *Pythium ultimum* were aligned using COBALT, and BOXSHADE was used to visualize conservation. Amino acid residues identical in all sequences are highlighted in black, while those identical in >80% of the sequences are highlighted in grey. (b) Percentages of amino acid sequence identity of the elicitin domain between the different elicitins. (c) A neighbor-joining phylogenetic tree was created based on the elicitin domains of the seven elicitins using MEGA 5.1. Bootstrap support values (1,000 replicates) above 50% are given next to the branches.

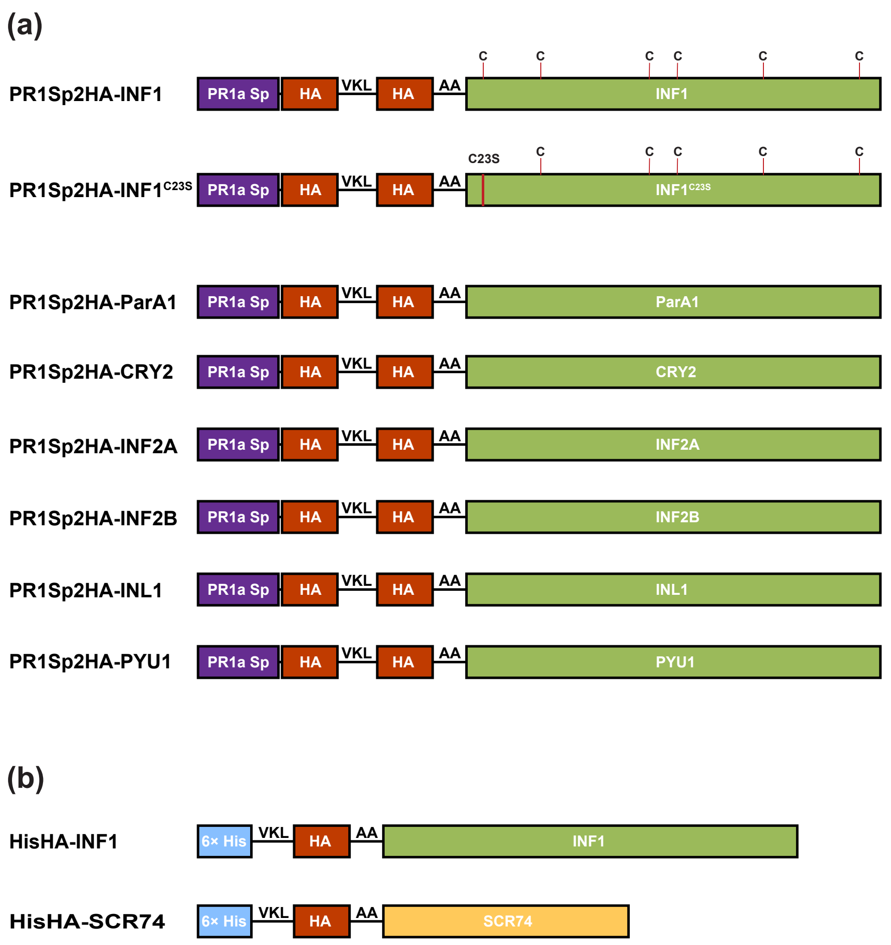


**Fig. S2. Overview of affinity-tagged elicitins used in this study.** Effector constructs used for (a) transient expression *in planta* using *Agrobacterium tumefaciens* and for (b) recombinant protein expression in *Pichia pastoris*. Pathogenesis-related protein 1a (PR1a) or native INF1 signal peptides (Sp) for extracellular targeting were used, while elicitins were carrying hemagglutinin (HA) or histidine (His) tags, as indicated. The 98-amino acid elicitin domains were used for all tested elicitins.

**Table S1**. Primers used in this study.

| Primer name | Purpose | Sequence (5′ to 3′) |
|---|---|---|
| PR1Sp Fwd | Cloning in pENTR/D-TOPO | CACCATGGGATTTGTTCTCTTTTCACAA |
| INF2A eli Rev | Cloning of elicitin domain[1] | TTACGACGAGGAGCACTTCTC |
| INF2B eli Rev | Cloning of elicitin domain[1] | TTACGACGAGGAGCACTTCTT |
| CRY2 eli Rev | Cloning of elicitin domain[1] | TTACAGCGAAGCACACGTC |
| INL1 eli Rev | Cloning of elicitin domain[1] | TTAAGCGCCAGCGCAGT |

[1] Jiang et al. (2006)
